# Effect of time spent on active learning on exam performance: A controlled case study on a course with different instructors but identical teaching materials

**DOI:** 10.1101/2022.09.01.506238

**Authors:** Xinjian Cen, Rachel J. Lee, Christopher Contreras, Melinda T. Owens, Jeffrey Maloy

## Abstract

Active learning, including student thinking and discussion in class, has been shown to increase student learning gains. However, it is less clear how variations in how instructors implement active learning affect student gains. Our study aims to investigate the extent to which the time spent on individual episodes of active learning activities influences student performance. We hypothesized that instructors who let students spend more time on peer discussion and individual thinking on practice problems associated with particular learning objectives will have better student exam scores on exam questions addressing those objectives. To test this hypothesis, we obtained a large data set of classroom recordings and student exam scores from an introductory biology course at a large four-year university, where three instructors shared identical teaching materials and exams for different course offerings.

Contrary to our hypothesis, although the three instructors spent significantly different amounts of time on episodes of thinking and peer discussion, there was no correlation between the total time spent on active learning activities and student performance on exam questions. Linear mixed-effects modeling of the effect of length of episodes of student thinking and discussion on exam score found that the amount of course time spent on active learning activities did not reliably predict student performance on associated exam questions. This result held true even when only considering learning objectives with high variations in performance between offerings, difficult exam questions, exam questions requiring higher-order thinking skills, or within-instructor performance. Although our study was only conducted in one course, our results imply that time spent per individual episode of student thinking or peer discussion may not be the primary factor explaining the positive effects of active learning and that it may be worthwhile to explore other factors.

## Introduction

Education in science, technology, engineering, and mathematics (STEM) is crucial for students entering careers necessary for national success [1]. Between 2000 and 2019, the number of STEM baccalaureate degrees awarded in the United States rose from about 400,000 to just under 725,000 [2]. Along with this increase there has been a renewed focus on committing instructors and institutions to adopt evidence-based teaching practices demonstrated to enhance students’ understanding of essential concepts in STEM [3,4]. Despite substantial progress in research on what types of teaching techniques most effectively improve student learning, there is still a gap in our understanding about the mechanistic underpinnings of these techniques [5]. As a result, there is considerable variability in what instructors who claim to use similar instructional practices actually do in the classroom, which may contribute to variable student outcomes [6]. Therefore, it has become vital to identify the mechanistic contributors to the efficacy of evidence-based instructional practices to improve the dissemination and implementation of effective instructional techniques in the classroom.

Many decades of research have shown that learning is more effective when learners actively construct knowledge by assimilating new information into their existing worldview rather than passively taking in information, a well-known framework known as constructivism [7]. “Active learning” refers to a diverse set of teaching strategies inspired by a constructivist framework; these practices involve students in doing things and thinking about what they are doing rather than simply being physically present in an educational environment [8]. Numerous studies have shown that when compared to traditional lectures, active learning environments increase student performance and decrease fail rates [9,10]. A meta-analysis by Theobald et al. further demonstrated that active learning not only benefits all students but especially helps to decrease grade gaps for minoritized students [11]. However, and it is not always clear what specific teaching practices produce these benefits, as active learning can be hard to precisely define [12,13]. For Auerbach and Schussler, active learning is defined as a student-centered pedagogy that incorporates student engagement and thinking through class activities where students collaborate and explain their understanding of concepts [14]. Others have defined active learning as the act of knowledge synthesis based on constructivism [15–18]. Still others define it as the absence of passive learning, with the goal of active learning being to minimize the time spent passively absorbing information taught through lecture [19–22]. Given such diverse definitions, it is not surprising that teaching strategies as disparate as cooperative learning, clicker questions, worksheets, interactive demonstrations, peer instruction, problem-based learning, and ability-based education are all called “active learning” in different contexts [23,24]. To amalgamate these different ideas about active learning, Driessen et al. propose a broad definition of active learning as, “an interactive and engaging process for students that may be implemented through the employment of strategies that involve metacognition, discussion, group work, formative assessment, practicing core competencies, live-action visuals, conceptual class design, worksheets, and/or games” [13].

Despite the differences between various active learning contexts, one commonality of many of the active learning activities and definitions described above is the presence of students thinking and discussing materials in class [25]. If students are thinking and discussing, they are not passively listening to the lecture. Instead, they are engaged, thinking, collaborating, and explaining, and at least some portion of class time is spent with the focus on the students instead of the instructor. Such activities can lead to active knowledge construction and the active assimilation of new information into students’ existing knowledge structures [26]. Because of the direct tie between student thought and discussion and constructivist ideas about effective learning, it is possible that they may underlie many of the theoretical mechanisms of action describing why active learning benefits students intellectually, motivationally, and socially. For example, when students ponder and explain material to each other in class, these actions may promote advanced levels of cognitive thinking, promoting their conceptual development and helping them uncover their own misconceptions [27,28]. Active thought and discussion may also stimulate student motivation and increases their engagement [29,30]. Finally, as students talk and form connections with each other, it could increase feelings of community and belonging [31,32].

Due in part to increasing professional development opportunities and an overwhelming body of literature describing the benefits of including student thought and discussion during class, instructors are increasingly aware of practices meant to promote these behaviors in class, and a large number of instructors claim to incorporate student thought and discussion at least occasionally in their classes [33–35]. However, observational studies have indicated a large degree of variability in practice [36]. In part, this variability may be explained by the lack of consensus on best practices for implementation of specific active learning techniques. When an instructor includes student thinking and discussion in their class, they need to make numerous decisions about implementation. Even for a basic clicker question, a commonly-used active learning activity where students use a clicker device to respond to multiple choice questions prompted by the instructor, the instructor needs to decide the difficulty and wording of the question, whether to force students to think about the question individually, whether to allow the students to discuss it with peers, how long students are given to think or discuss, and how to grade responses [36–38]. Different decisions in any of these aspects of implementation may affect how well instructors assess student understanding, provide student-centered feedback, and ultimately help students learn the material.

One understudied aspect of active learning implementation is the amount of time an instructor should spend on individual episodes of student thinking and discussion. In general, the evidence points to a positive correlation between the time spent on active learning and better student outcomes. Elliott et al. and Preszler et al. found that courses with more clicker questions generally had higher student grades, and Moon et al. found that greater total time spent in student thought, work, or talk correlated with greater exam scores [39,40]. Theobald et al. also showed that grade gaps between minoritized and non-minoritized students are the smallest in the courses with the most active learning [11]. For this reason, much informal advice encourages instructors to increase the amount of classroom time spent in active learning [e.g, 41]. However, these studies do not necessarily imply that instructors should strive to lengthen individual episodes of student thinking and discussion. These studies compared different courses, which were also likely to be different in various confounding factors that may affect student learning and achievement, including the number and type of active learning activities as well as other factors like learning objectives, course content, lecture materials, and assessment methods. Studies that have specifically looked at the length of time spent on individual episodes of student thinking and discussion have found that although instructors vary in the amount of time they give students to think about, discuss, and report their ideas, the effect of such differences on student learning is unclear [42].

In this study, we controlled for these confounding factors by analyzing a model system where all teaching and assessment materials were identical across different offerings, so time spent on student thinking and discussion was one of the only active learning variables that instructors could alter. With this model, we asked whether the time spent on student thinking and discussion correlated with increased student learning as measured by performance on course exams. We hypothesized that instructors who allow students to spend more time on silent thinking and peer discussion associated with particular learning objectives would have better student exam scores on exam questions addressing those objectives.

## Materials and Methods

### Study Context

The study utilized a large data set of classroom video recordings and student exam scores from an introductory biology course at a large, public, West Coast R1 university. Permission for this study was obtained from the UCLA Human Research Protection Program in certified exempt protocol #19-001554. The course was in-person and made extensive use of active learning. Students in the course were assigned pre-class readings and recorded content videos to be completed before class. Their pre-class comprehension was verified using pre-class review quizzes and the submission of pre-class reading guides. During class, the active learning activities in this course consisted of applying the knowledge obtained from pre-class activities to novel problems using in-class clicker questions, peer discussion questions, and worksheet questions. Typically, the instructors would direct the students to work on the clicker questions or discussion activities with their peers, although occasionally, students were asked to think individually or given no instructions. A certain amount of time was then allotted for students to complete the activity before the instructor regrouped and discussed the question with the students. Roughly 15-20% of class time was spent in student individual thought or peer discussion, as measured by the techniques outlined in the “Analysis of time spent on active learning” section below (Supplemental Figure 1). It should be noted that this percentage does not include the time the instructor took to explain the question or activities or to follow up on the questions, so the total class time devoted to active learning activities is much higher.

In the specific quarter studied, the course had three offerings of 340-380 students each, with each offering taught by a different instructor. All instructors were full-time non-tenure-track instructors with multiple years of experience teaching and coordinating the course who have received professional development on implementing active learning. Each instructor used identical teaching materials, including Powerpoint slides, clicker questions, and worksheets. They also all used the same comprehensive list of learning objectives, which corresponded to each class unit and aligned with exam questions. They worked together to develop the course teaching materials and agreed to not deviate from them, which was informally confirmed through viewing the video recordings of their classes. The assessments were also all identical and jointly approved. The course had two midterms and a final exam, all in-person. Exams were solely multiple-choice to remove subjectivity in grading. All students answered the same questions, although there were two different versions of the exam with the questions shuffled to minimize cheating. Thus, student performance is directly comparable across the course offerings.

### Assessment of prior knowledge

All three offerings of the course analyzed in this study were listed on the registrar’s course enrollment website simultaneously with identical course descriptions. To ensure that the beginning student populations in each offering were comparable, students were administered the General Biology Measuring Achievement and Progression in Science (GenBIO-MAPS) concept inventory assessment (Supplemental Figure 2) during the first week of the course [43]. All students were asked to attempt the assessment for two bonus points in the course. Following this assessment period, we retrieved student results and filtered to remove any students who did not meet the following criteria:

a. Completed the entire assessment
b. Confirmed that they were at least 18 years of age
c. Took at least 10 minutes to complete the assessment
d. Remained enrolled in the course at least through the administration of the first exam

Overall, assessment completion rates were under 50% for all instructors (instructor A: 35.70% (146/409), instructor B: 48.52% (197/406), instructor C: 42.09% (181/430).

### Choosing teaching units to analyze

Because of the volume of exam and active learning questions, we initially chose specific teaching units to analyze that had the highest standard deviation of performance between instructors. Each teaching unit covered between three and seven learning objectives. One teaching unit was chosen from each third of the course, so that one unit each was tested on Midterm 1, Midterm 2, and the Final Exam. The number of observations, clicker questions, and exam questions analyzed per teaching unit are shown in Supplemental Table 1.

Later, we analyzed questions that were difficult. For this analysis, we analyzed all learning objectives associated with exam questions where under 56% of students received credit for a correct answer. This cutoff was chosen empirically, because we saw a question where there was a possible correlation between length of discussion time and student score.

### Classification of cognitive order for clicker and exam questions

To further our analysis of question type difficulty, our team classified the cognitive complexity of clicker questions and exam questions into higher cognitive order (higher-order) and lower cognitive order (lower-order) question using Bloom’s Taxonomy following previously published guidelines [44,45]. Higher-order questions encompassed Bloom’s *analyze*, *create*, and *evaluate*, while lower-order questions consisted of Bloom’s *apply, remember*, and *understand*. According to the Larsen et al. 2022 definition of *apply*, we classified *apply* under lower-order because the *apply* questions in this course fit the subcategory *execute*, when students were expected to follow a procedure for a similar problem that was learned during class time. Sample lower- and high-order clicker and exam questions from the course can be found in Supplemental Figure 3. Two or three members of the research team Bloomed each question, and if there was disagreement, determinations were made through consensus among the research team. Since some learning objectives were tested using both higher and lower-order questions, we calculated percent of higher and lower-order questions exam questions for each learning objective (Supplemental Table 2).

### Analyzing exam performance

Exam questions were grouped by learning objective before analysis. Researchers matched all of the exam questions and classroom active learning activities to their corresponding learning objectives. To find the exam performance for a particular learning objective for a particular instructor, we averaged that instructor’s students’ scores (average correctness) for all the exam questions addressing that learning objective. To compare performance within instructors, the exam scores for each learning objective were normalized to reflect exam performance of a specific instructor relative to the average exam performance of all three instructors on that objective. This was done by subtracting the aggregate average score of all three instructors from the average score of that instructor’s students.

### Analysis of time spent on active learning

Time spent on active learning was calculated from video recordings. These video recordings were not recorded for research purposes; instead, they were automatically generated from cameras positioned in the back of classrooms to help students review lectures. Thus, they only captured instructor and screen activity in the front of the classroom. Audio was recorded through the instructors’ lapel microphones. Instructors often turned the microphones and therefore the audio-recording off while students were actively thinking or discussing.

To measure time spent in active learning, two researchers hand-coded the length of each active learning opportunity from lecture recordings. More specifically, they used slide transitions and instructor cues to mark the times each clicker question or classroom activity opened and closed (Supplemental Figure 4). The time spent on each episode of student thinking or discussion was calculated from the timestamps. We did not distinguish between episodes of student thinking and student discussion because in this course, instructors often did not ask for only one of these to occur, so it was likely that both occurred in overlapping time periods. The time spent on active learning per learning objective for each instructor was calculated by summing the total time spent on active learning activities (in seconds) for a particular instructor for each question that addressed the learning objective. To address inter-rater reliability, 10% of the active learning episode was coded independently by two researchers. Afterwards, the researchers compared their timestamps and resolved any discrepancies. If the coders disagreed on the timing of any activity’s open or close time, the researchers discussed their decisions and tried again with another 10% of the active learning episodes until the coders could consistently, 100% of the time independently choose timestamps within 5 seconds of each other. The remaining videos were coded independently by one researcher.

### Statistical analysis

To determine whether the three instructors spent different lengths of time on student thinking or discussion and to determine whether exam scores were different between the instructors, we obtained the p-value from a randomization test using the PlotsOfDifferences tool, to which we then applied a Bonferroni correction, adjusting the significance threshold by a factor of 3 to p<0.0125 [46]. Randomization tests are a way of determining the significance of differences between groups that do not rely on assumptions about the distributions of the data [47]. To determine whether there was a significant correlation between student performance and time spent in student thinking and discussion, we developed a linear mixed-effects model using the “lmer” command in the “lme4” package in R [48]. To account for random effects of different instructors and learning objectives, we used the following model for all analyses in this study: average score ∼ length + (1|instructor) + (1|objective). Statistical test results were generated with the “LmerTest” package, including corrected p-values and summary tables for lmer model fits via Satterthwaite’s degrees of freedom method (Table 1) [49].

**Table 1.**
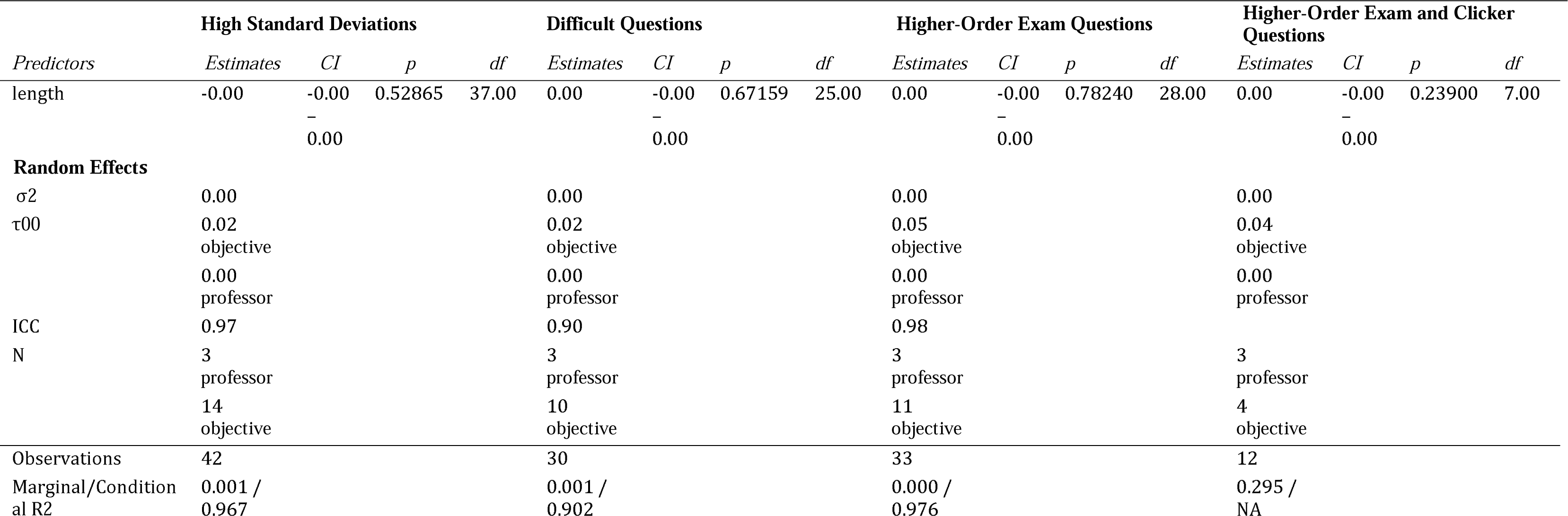
Linear Mixed Effects Model Results.

| *Predictors* | High Standard Deviations |  |  |  | Difficult Questions |  |  |  | Higher-Order Exam Questions |  |  |  | Higher-Order Exam and Clicker Questions |  |  |  |
| --- | --- | --- | --- | --- | --- | --- | --- | --- | --- | --- | --- | --- | --- | --- | --- | --- |
|  | *Estimates* | *CI* | *p* | *df* | *Estimates* | *CI* | *p* | *df* | *Estimates* | *CI* | *p* | *df* | *Estimates* | *CI* | *p* | *df* |
| length | -0.00 | -0.00<br>-<br>0.00 | 0.52865 | 37.00 | 0.00 | -0.00<br>-<br>0.00 | 0.67159 | 25.00 | 0.00 | -0.00<br>-<br>0.00 | 0.78240 | 28.00 | 0.00 | -0.00<br>-<br>0.00 | 0.23900 | 7.00 |
| **Random Effects** |  |  |  |  |  |  |  |  |  |  |  |  |  |  |  |  |
| σ<sup>2</sup> | 0.00 |  |  |  | 0.00 |  |  |  | 0.00 |  |  |  | 0.00 |  |  |  |
| τ<sub>00</sub> | 0.02 |  |  |  | 0.02 |  |  |  | 0.05 |  |  |  | 0.04 |  |  |  |
|  | objective |  |  |  | objective |  |  |  | objective |  |  |  | objective |  |  |  |
|  | 0.00 |  |  |  | 0.00 |  |  |  | 0.00 |  |  |  | 0.00 |  |  |  |
|  | professor |  |  |  | professor |  |  |  | professor |  |  |  | professor |  |  |  |
| ICC | 0.97 |  |  |  | 0.90 |  |  |  | 0.98 |  |  |  |  |  |  |  |
| N | 3 |  |  |  | 3 |  |  |  | 3 |  |  |  | 3 |  |  |  |
|  | professor |  |  |  | professor |  |  |  | professor |  |  |  | professor |  |  |  |
|  | 14 |  |  |  | 10 |  |  |  | 11 |  |  |  | 4 |  |  |  |
|  | objective |  |  |  | objective |  |  |  | objective |  |  |  | objective |  |  |  |
| Observations | 42 |  |  |  | 30 |  |  |  | 33 |  |  |  | 12 |  |  |  |
| Marginal/Conditional R<sup>2</sup> | 0.001 / 0.967 |  |  |  | 0.001 / 0.902 |  |  |  | 0.000 / 0.976 |  |  |  | 0.295 / NA |  |  |  |

## Results

Because the three instructors observed in this study used identical course materials including clicker questions and active learning segments, we might expect that all instructors spent similar amounts of time on student thinking and discussion. However, statistical analysis via a randomization test shows a significant difference in the lengths of active learning episodes per instructor, with instructor B having much longer active learning episodes than instructor A (p<0.001; Cohen’s d: 0.827; effect size: 45.593 seconds), who had moderately longer active learning episodes than instructor C (p=0.048; Cohen’s d: 0.433; effect size: 16.159 seconds) (Fig 1A).

**Fig 1.**
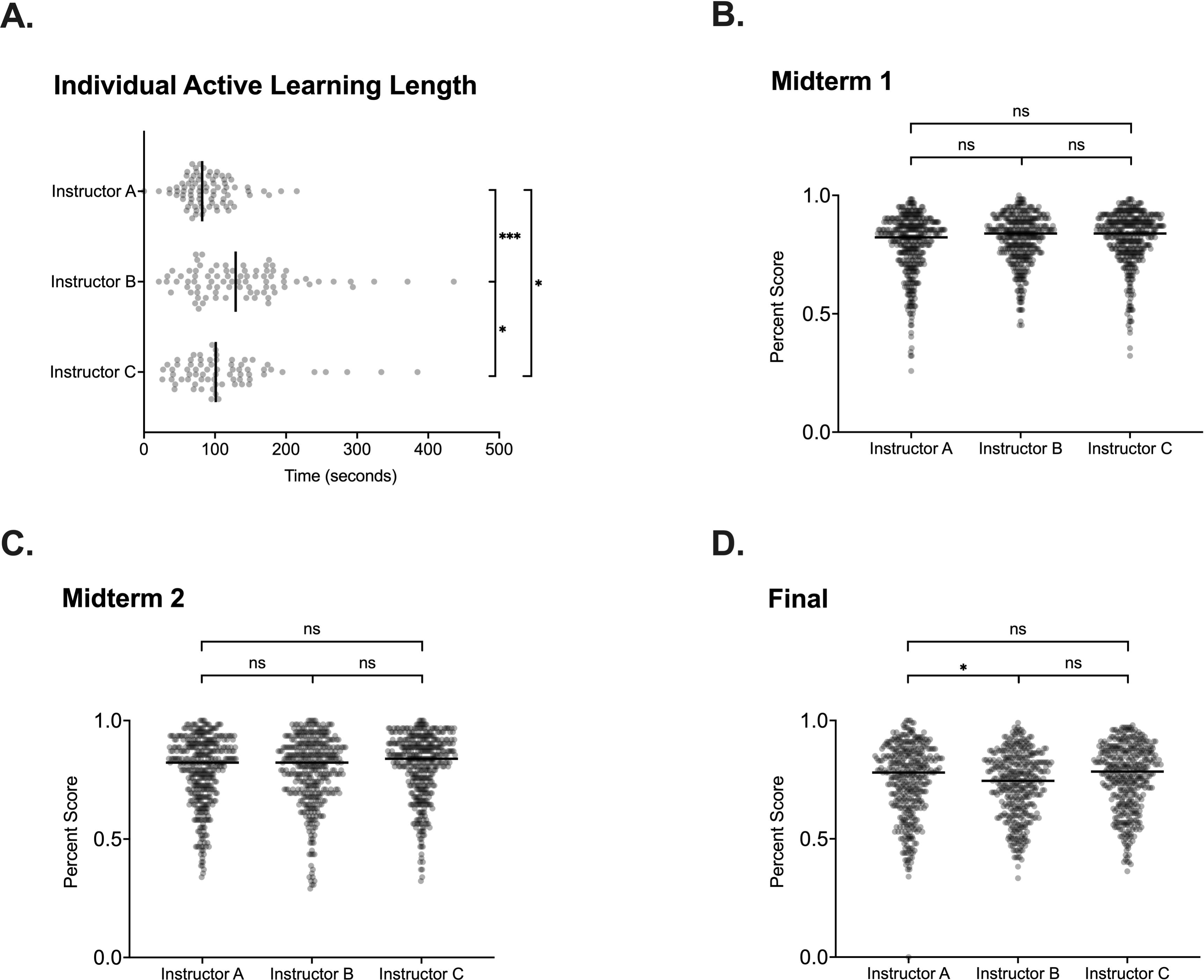
Overall trends in lengths and scores among the instructors. (A) Length of the individual active learning episode lengths for each instructor. (B-D) Percent scores of all individual students on midterm 1, midterm 2, and the final exam, respectively. * p≤0.0125, ns p>0.0125

Based on previous literature indicating that more active learning leads to better outcomes for students, with these differences in the amount of student thinking and discussion, one might also expect large differences in exam performance between each instructor’s students [11]. However, there were no significant differences in the overall exam scores for any instructor on either midterm. For the final exam, instructor A, who had the shortest average episode lengths, had slightly higher scores than instructor B (p=0.012; Cohen’s d: 0.112; effect size: -2.69%) (Figs 1B-D).

We do not believe that differences in the pre-course knowledge or preparation of students influence our results. In the first week of class, student knowledge was measured with the General Biology Measuring Achievement and Progression in Science (GenBIO-MAPS) concept inventory. Using a randomization test, we observed no significant differences between each instructor’s beginning student populations’ assessment percentage scores (all p>0.0125) (Supplemental Figure 2).

Because overall exam performance was similar between course offerings, to address our question of whether increased active learning time contributes to improved student outcomes, we categorized the exams questions into discrete learning objective units and analyzed these units separately. Since students in each course offering performed similarly on many of the learning objectives on the exams, we first decided to focus on the learning objectives with high variability in performance between course offerings. Using linear mixed modeling, for these learning objectives, we observed no correlation between time spent on student thinking and discussion and exam scores (p=0.5299) (Fig 2A, Table 1). However, we noticed that the two learning objectives in the high standard deviation sample that were particularly difficult for students, as indicated by average correctness of ≤56%, did show a strong correlation between exam score and time spent on student thinking and discussion, as seen by the gold and black lines in Fig 2A (r2=0.948 (black), r2=0.523 (gold)).

**Fig 2.**
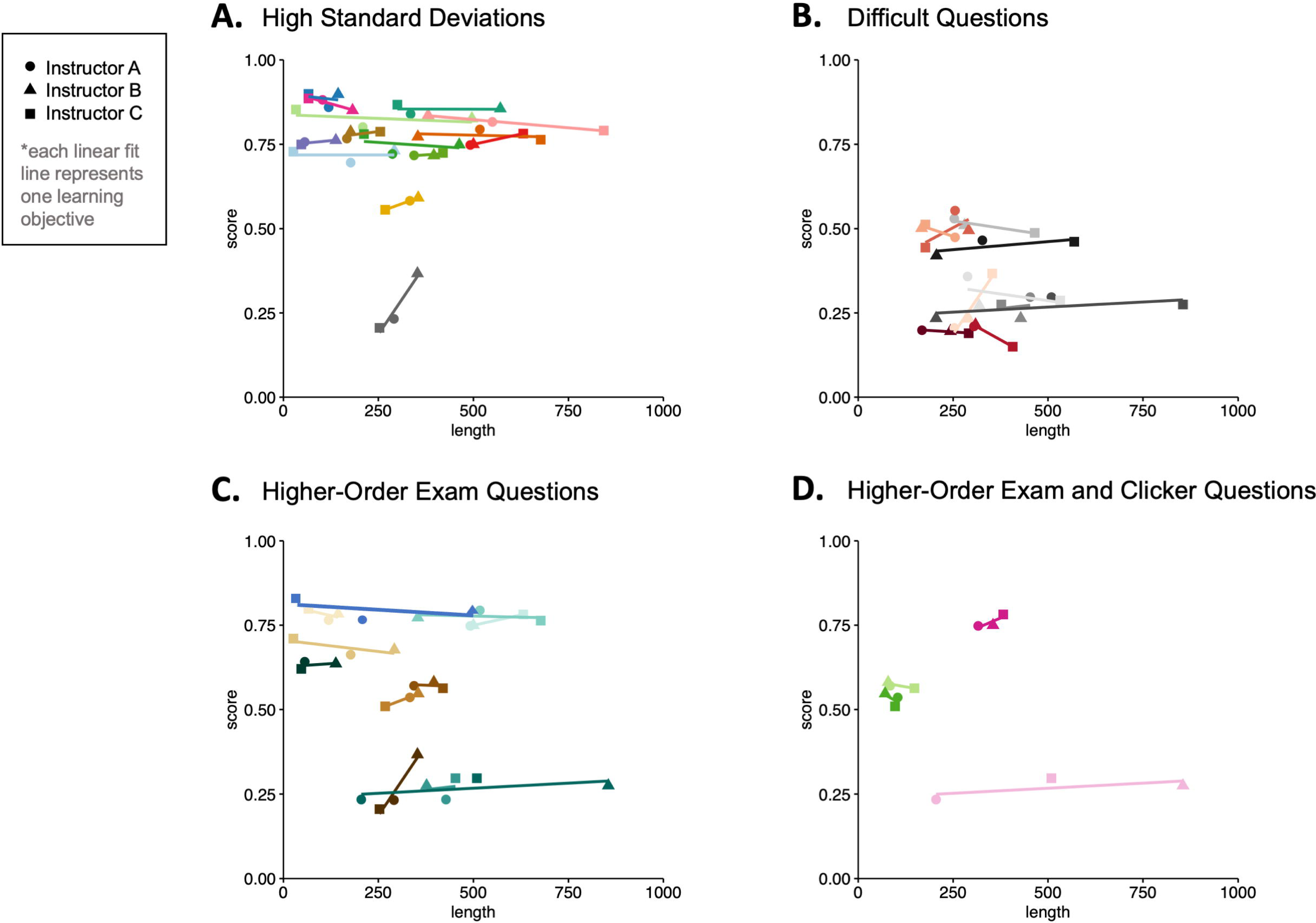
Effect of length of student discussion on exam score. Each line represents one learning objective. Within each panel, different colors represent different learning objectives. (A) Exam questions with high variability between instructors (p=0.5299). (B) Exam questions with <56% average correctness (p=0.6741). (C) Higher-order exam questions (p=0.7824). (D) Higher-order exam questions that have corresponding higher-order clicker questions (p=0.2390).

Given this observation, we hypothesized that increased time spent on student thinking and discussion might be beneficial for students for understanding and practicing difficult learning objectives. Thus, there might be a correlation between exam score and time spent on student thinking and discussing for a subset of learning objectives corresponding to very difficult exam questions. Thus, we applied linear mixed modeling to study teaching units that contained learning objectives corresponding to difficult exam questions, which we defined as questions with average correctness below 56%, which corresponds to the highest average score where we saw a possible correlation in our previous analysis. We still found no correlation between the length of time spent on active learning and student performance on difficult exam questions (p=0.67) (Fig 2B, Table 1).

While average correctness is a possible indicator of the difficulty level of a question, there is also the possibility that a question can have lower average scores because it is worded poorly or has a confusing sentence structure. Thus, to conduct a more in-depth analysis, we considered another category of difficult questions: those that require higher-order cognitive skills as determined by Bloom’s Taxonomy [44,45]. Such an approach has been used in the past to correlate exam difficulty with student performance [39,50]. However, in our sample, we determined that there was no correlation between the length of time spent on active learning and student performance on higher-order exam questions (p=0.78) (Fig 2C, Table 1) and on higher-order exam questions that have corresponding higher-order clicker questions (p=0.24) (Fig 2D, Table 1).

Finally, we investigated the possibility that within a given instructor, topics where students were given more time to think and discuss might show increased learning versus other topics taught by the same instructor with less active learning time. In other words, given that the class only lasted a fixed amount of time, perhaps an instructor’s students learned better on concepts where that instructor happened to allow the students to spend more time thinking and discussing. Therefore, we correlated the effects of time spent on active learning and student exam performance within each instructor for all the learning objectives, normalized for each exam’s average score. None of the instructors showed a significant correlation (Fig 3).

**Fig 3.**
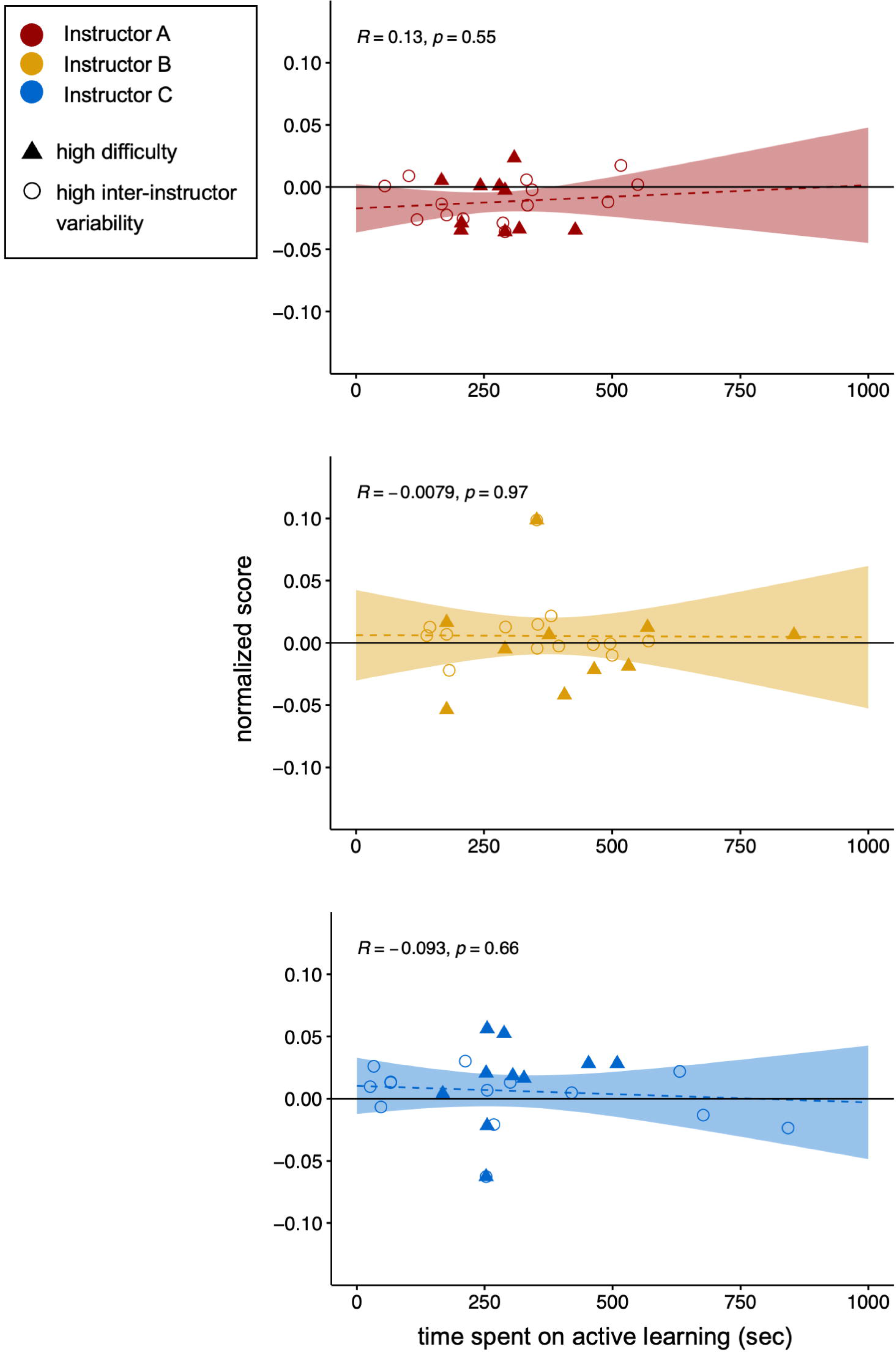
Normalized lines of regression per instructor. (normalized score v. time spent on active learning) (95% confidence intervals are shown as shaded areas and p-values are labeled on each plot). A data point above the x-axis indicates that an instructor’s exam score on a particular learning objective is above the average exam score for a particular learning objective across all three instructors.

## Discussion

Our case study is a controlled observational study in which different offerings of the same course had different instructors but the same set of teaching materials. This setup meant that instructors could only vary in how they implemented that teaching material, one aspect of which is the amount of time spent on student discussion and thinking. Overall, somewhat surprisingly, we found no effects of spending more time on student thinking and discussion on exam performance. Although the three instructors gave substantially different amounts of time to student thinking and discussing (Fig 1A), that variation was not reflected in student exam performance. In fact, instructor A had higher Final Exam scores than instructor B despite giving students shorter amounts of time to think and discuss (Fig 1D). There was no relationship between exam performance and time on student thinking and discussion even when only considering learning objectives with high variations in performance between offerings (Fig 2A), difficult exam questions (Fig 2B), exam questions requiring higher-order thinking skills (Fig 2C and 2D), or within-instructor performance (Fig 3).

This result was unexpected and is difficult to explain. Constructivism as well as a host of individual studies show that active learning, which includes student thinking and discussion, generally promotes student learning [9]. A previous study documented that in a similar system where two instructors shared common instructional materials, the instructor who spent more class time on having students discuss questions also had higher exam scores [51]. However, those two instructors had much larger differences in how much time they spent in student discussion (roughly 5% compared to roughly 30% of class time) than we saw, which may account for the larger differences they saw in student outcomes. Another study that also looked at time spent on student thinking or discussion found it had a small but significant positive effect on exam scores, but this study looked across courses, which introduces other confounding factors like course content and difficulty [39].

## Implications for Education

One possible way to reconcile our results with existing literature is to separate the effects of giving students more time on fewer clicker questions as opposed to giving students more questions. For example, Preszler et al (2007) Elliott et al (2016) did find a correlation between student learning gains and classroom time spent in active learning [40,52] However, in the context of those studies, greater classroom time spent on active learning arose from including more clicker questions, rather than increasing student discussion time for a constant number of clicker questions. Moon et al. (2021) found separate, positive effects for increasing time spent on student thinking and discussion and for the number of active learning activities when comparing different courses [39]. Therefore, for instructors trying to determine how much of their limited classroom time to devote to peer discussion, our data combined with previous literature suggests that increasing the number of clicker questions and active learning activities may be a more reliable way to reap the benefits of active learning, rather than giving more time to a small number of activities.

Our study also adds to the literature about educational “fidelity of implementation,” which has implications for curriculum development [53]. We found that even when instructors were given identical teaching materials, instructors implemented them differently (Fig 1A). That also accords with the results of Lo et al., who found that two instructors who co-created and used identical teaching materials spent significantly different amounts of time allowing students to discuss questions, answering student questions, and personally stimulating and supporting student discussion [51]. Many groups have tried to create instructional change by disseminating curricula, but there is significant controversy over whether these efforts are effective, in part due to instructor variability in implementation [53,54]. Our study’s results suggest that although instructors do indeed make their own decisions regarding implementation and variability is introduced between instructors when they are using co-constructed materials, this variability may not necessarily lead to different outcomes for students.

Finally, our study suggests that even though the link between active learning and better student outcomes is so strong, we cannot blindly assume that every instance of more student thinking and discussion automatically equates to better learning. This caution is especially important given that there are several established classroom observation tools that measure the amount and percentage of class time spent on various active learning activities like peer discussion, which implies that the time spent in active learning can give important information about teaching. Examples include the Classroom Observation Protocol for Undergraduate STEM (COPUS), which uses a set of codes to categorize student and instructor behavior within every two minutes of class time [55], Decibel Analysis for Research in Teaching (DART), which analyzes the volume and variance of classroom audio-recordings to estimate the percentage of time spent in active learning [56], and the Practical Observation Rubric To Assess Active Learning (PORTAAL), which measures the minutes students could spend in thinking or discussion as a component of its score [57]. Although the creators of such protocols are clear in stating measuring time spent on classroom activities cannot tell you about the quality and therefore the effect of that instruction, there has been a tendency among users to over-interpret observation protocol findings and assume that teaching styles with more time spent on active learning are inexorably better [41,58,59]. While it is clear that pure lecturing typically produces poor learning compared to many kinds of active learning, there are many other factors that influence teaching effectiveness and it matters how that active learning is implemented [9,58]. Our study adds to the literature showing that “more time” may not necessarily mean “better” and that although protocols like COPUS and DART give useful information, their results should ideally be put in greater context especially in high-stakes situations like instructor evaluations.

## Limitations

The nature of this study presents several limitations. One is that considerable amount of class time was already spent on students doing active learning, so the differences in time spent on student thinking and discussion between offerings may not have had a large effect. It is true that the differences in the time spent time on student thinking and discussion were significant with large to moderate effect sizes based on Cohen’s d. However, it is possible that increased student outcomes due to active learning were already “baked in” to the course design and that any additional variation in time spent on active learning by specific instructors produced diminishing returns compared to the effect of the overall course structure.

The fact that this study analyzed real courses also meant that there were many factors that could not be controlled for. The course offerings were held at different times of day, which may have affected student attendance or engagement with the learning activities. Students likely varied in how much time they spent and what techniques they used to study the course material outside of class. In addition, we did not analyze teaching techniques used during graduate TA-led discussion offerings or out-of-class exam review sessions. However, given that we did not see large differences between instructors in exam scores, it is likely that these factors did not lead to systematic differences between course offerings.

We also did not consider the quality of student interactions during active learning. The audio recorders were typically turned off during active learning activities, preventing researchers from observing or hearing student activity. This raises questions about how engaged students were, as some of them may have been doing off-topic activities during the thinking and discussion period.

Lastly, this is a case study, and the curriculum and setting of the course were specific to the university of interest. We also recognize that small class settings, such as discussion sections, offer a different dynamic of active learning in the instructor-student and student-student interaction. Thus, findings may not generalize to other courses. More work will be needed to determine whether there may be benefits of student thought and discussion time that are context-specific.

## Future directions

There are many possible future directions to this study. One factor that can be explored would be the type of behavior and discourse that students were engaged in during the activ learning time period. For instance, audio recording students during active learning would allow researchers to code their behaviors (whether they are engaged in peer discussion, individual thinking, or studying independently) and discourse patterns. Studies have shown that students who chose to engage in peer discussion/instruction showed better learning outcomes than students who completed activities individually [60,61]. Then, instructors could better understand the connections between how they implement active learning, how students respond and carry out the active learning, and student outcomes.

Also, although we found that different instructors had differences in discussion timing, it is not clear why such variations arise. Are these differences coming about because instructors are attending to different classroom cues, like the amount of noise; doing different activities like spending time with individual student groups during the discussion period; or making different decisions about what percentage of the students have submitted answers [42,51]? so, how do such differences relate to differences in instructor ideas about learning, teaching and diversity? Understanding why instructors make implementation decisions will greatly benefit faculty development efforts.

## Conclusion

Overall, our results imply that simply varying the amount of time spent on given active learning activities may not contribute to differential student learning gains. It would be worthwhile to further explore other variables that might enhance the positive effects of activ learning, such as number or variety of clicker questions or activities or how instructors support or follow up on student discussions. Additionally, we show that student learning is robust to some differences in implementation of active learning techniques, suggesting that dissemination of shared teaching materials between instructors with similar levels of trainin and experience with active learning techniques may be an effective means of reaping the benefits of active learning in more classrooms. Moving forward, similar studies using co-constructed and shared materials with controlled deviations per instructor may help to discern the possible mechanisms by which active learning activities impact students in the undergraduate classroom.

## Acknowledgements

We are grateful for the instructors and students whose efforts we studied. We would also like to thank the UCLA and UCSD biology education research communities for helpful feedback.

## Supporting information

S1 Table. Description of class sessions and questions analyzed in this study.

S1 Figure. Proportion of class time spent on active learning activities relative to non-active learning activities.

S2 Figure. GenBIO-MAPS concept inventory scores across instructors

S3 Figure. Selected examples of higher- and lower-order clicker and exam questions analyzed

S4 Figure. Guidelines used by coders for how to determine starting and ending times for each active learning activity.

## Notes

### Competing Interest Statement

The authors have declared no competing interest.

### Summary of Updates

We have added an additional analysis comparing class performance on a pre-test before the start of the term. We have also incorporated analyses of Bloom's levels for different clicker and exam questions analyzed, to see if results varied depending on whether questions were low- or high- cognitive order. We have also revised the introduction and discussion of the paper. An additional author, Christopher Contreras, was involved in the new analyses and has been added to the author list.

## References

1. Olson S, Riordan DG. Engage to Excel: Producing One Million Additional College Graduates with Degrees in Science, Technology, Engineering, and Mathematics. Report to the President. Executive Office of the President. 2012.

2. Trapani J, Hale K. Higher Education in Science and Engineering. Science & Engineering Indicators 2022. NSB-2022-3. National Science Foundation. 2022.

3. Singer SR, Nielsen NR, Schweingruber HA. Discipline-based education research. Washington, DC: The National Academies. 2012.

4. Education NSF (US) D for, Resources H. Shaping the future: New expectations for undergraduate education in science, mathematics, engineering, and technology. National Science Foundation, Division of Undergraduate Education; 1996.

5. Streveler RA, Menekse M. Taking a Closer Look at Active Learning. J Eng Educ. 2017;10 186–190. doi:10.1002/jee.20160

6. Stains M, Harshman J, Barker MK, Chasteen SV, Cole R, DeChenne-Peters SE, et al. Anatomy of STEM teaching in North American universities. Science. 2018;359: 1468– 1470. doi:10.1126/science.aap8892

7. Driver R, Asoko H, Leach J, Scott P, Mortimer E. Constructing scientific knowledge in th classroom. Educational researcher. 1994;23: 5–12.

8. Bonwell CC, Eison JA. Active learning: Creating excitement in the classroom. 1991 ASHE-ERIC higher education reports. ERIC; 1991.

9. Freeman S, Eddy SL, McDonough M, Smith MK, Okoroafor N, Jordt H, et al. Active learning increases student performance in science, engineering, and mathematics. PNAS. 2014;111: 8410–8415. doi:10.1073/pnas.1319030111

10. Dewsbury BM, Swanson HJ, Moseman-Valtierra S, Caulkins J. Inclusive and active pedagogies reduce academic outcome gaps and improve long-term performance. Plos one. 2022;17: e0268620.

11. Theobald EJ, Hill MJ, Tran E, Agrawal S, Arroyo EN, Behling S, et al. Active learning narrows achievement gaps for underrepresented students in undergraduate science, technology, engineering, and math. Proceedings of the National Academy of Sciences. 2020;117: 6476–6483.

12. Dewsbury BM. Context determines strategies for ‘activating’the inclusive classroom. Journal of Microbiology & Biology Education. 2017;18: 18.3. 30.

13. Driessen EP, Knight JK, Smith MK, Ballen CJ. Demystifying the meaning of active learning in postsecondary biology education. CBE—Life Sciences Education. 2020;19: ar52.

14. Auerbach AJ, Schussler EE. Curriculum alignment with Vision and Change improves student scientific literacy. CBE—Life Sciences Education. 2017;16: ar29.

15. Bentley M, Connaughton VP. A simple way for students to visualize cellular respiration adapting the board game MousetrapTM to model complexity. 2021.

16. Connell GL, Donovan DA, Chambers TG. Increasing the use of student-centered pedagogies from moderate to high improves student learning and attitudes about biology. CBE—Life Sciences Education. 2016;15: ar3.

17. Cooper KM, Ashley M, Brownell SE. A bridge to active learning: A summer bridge program helps students maximize their active-learning experiences and the active-learning experiences of others. CBE—Life Sciences Education. 2017;16: ar17.

18. Stoltzfus JR, Libarkin J. Does the room matter? Active learning in traditional and enhanced lecture spaces. CBE—Life Sciences Education. 2016;15: ar68.

19. Hoefnagels M, Taylor MS. “ Boost your evolution IQ”: An evolution misconceptions gam 2021.

20. McLean JL, Suchman EL. Using magnets and classroom flipping to promote student engagement and learning about protein translation in a large microbiology class. Journal of microbiology & biology education. 2016;17: 288–289.

21. McCourt JS, Andrews TC, Knight JK, Merrill JE, Nehm RH, Pelletreau KN, et al. What motivates biology instructors to engage and persist in teaching professional development? CBE—Life Sciences Education. 2017;16: ar54.

22. Lee CJ, Toven-Lindsey B, Shapiro C, Soh M, Mazrouee S, Levis-Fitzgerald M, et al. Error-discovery learning boosts student engagement and performance, while reducing student attrition in a bioinformatics course. CBE—Life Sciences Education. 2018;17: ar40.

23. AlRuthia Y, Alhawas S, Alodaibi F, Almutairi L, Algasem R, Alrabiah HK, et al. The use of active learning strategies in healthcare colleges in the Middle East. BMC medical education. 2019;19: 1–10.

24. Mazur E, Hilborn RC. Peer instruction: A user’s manual. Physics Today. 1997;50: 68.

25. Linton DL, Pangle WM, Wyatt KH, Powell KN, Sherwood RE. Identifying Key Features o Effective Active Learning: The Effects of Writing and Peer Discussion. Momsen J, editor LSE. 2014;13: 469–477. doi:10.1187/cbe.13-12-0242

26. Michael J. Where’s the evidence that active learning works? Advances in physiology education. 2006.

27. Linnenbrink EA, Pintrich PR. Role of affect in cognitive processing in academic context Motivation, emotion, and cognition. Routledge; 2004. pp. 71–102.

28. Akınoğlu O, Tandoğan RÖ. The effects of problem-based active learning in science education on students’ academic achievement, attitude and concept learning. Eurasia journal of mathematics, science and technology education. 2007;3: 71–81.

29. Cicuto CAT, Torres BB. Implementing an active learning environment to influence students’ motivation in biochemistry. Journal of Chemical Education. 2016;93: 1020– 1026.

30. Armbruster P, Patel M, Johnson E, Weiss M. Active learning and student-centered pedagogy improve student attitudes and performance in introductory biology. CBE— Life Sciences Education. 2009;8: 203–213.

31. Meeuwisse M, Severiens SE, Born MP. Learning environment, interaction, sense of belonging and study success in ethnically diverse student groups. Research in Higher Education. 2010;51: 528–545.

32. Masika R, Jones J. Building student belonging and engagement: Insights into higher education students’ experiences of participating and learning together. Teaching in higher education. 2016;21: 138–150.

33. Borrego M, Froyd JE, Hall TS. Diffusion of engineering education innovations: A survey of awareness and adoption rates in US engineering departments. Journal of Engineerin Education. 2010;99: 185–207.

34. Prince M, Borrego M, Henderson C, Cutler S, Froyd J. Use of research-based instruction strategies in core chemical engineering courses. Chemical Engineering Education. 2013;47: 27–37.

35. Macdonald RH, Manduca CA, Mogk DW, Tewksbury BJ. Teaching methods in undergraduate geoscience courses: Results of the 2004 On the Cutting Edge survey of US faculty. Journal of Geoscience Education. 2005;53: 237–252.

36. Vickrey T, Rosploch K, Rahmanian R, Pilarz M, Stains M. Research-Based Implementation of Peer Instruction: A Literature Review. Barnard D, editor. LSE. 2015;14: es3. doi:10.1187/cbe.14-11-0198

37. Martyn M. Clickers in the classroom: An active learning approach. Educause quarterly. 2007;30: 71.

38. Ding L, Reay NW, Lee A, Bao L. Are we asking the right questions? Validating clicker question sequences by student interviews. American Journal of Physics. 2009;77: 643– 650.

39. Moon S, Jackson MA, Doherty JH, Wenderoth MP. Evidence-based teaching practices correlate with increased exam performance in biology. PLOS ONE. 2021;16: e0260789 doi:10.1371/journal.pone.0260789

40. Preszler RW, Dawe A, Shuster CB, Shuster M. Assessment of the Effects of Student Response Systems on Student Learning and Attitudes over a Broad Range of Biology Courses. CBE Life Sci Educ. 2007;6: 29–41. doi:10.1187/cbe.06-09-0190

41. Reisner BA, Pate CL, Kinkaid MM, Paunovic DM, Pratt JM, Stewart JL, et al. I’ve Been Given COPUS (Classroom Observation Protocol for Undergraduate STEM) Data on My Chemistry Class… Now What? J Chem Educ. 2020;97: 1181–1189. doi:10.1021/acs.jchemed.9b01066

42. Turpen C, Finkelstein ND. Not all interactive engagement is the same: variations in physics professors’ implementation of peer instruction. Physical Review Special Topic Physics Education Research. 2009;5: 020101.

43. Couch BA, Wright CD, Freeman S, Knight JK, Semsar K, Smith MK, et al. GenBio-MAPS: A programmatic assessment to measure student understanding of vision and change cor concepts across general biology programs. CBE—Life Sciences Education. 2019;18: ar1

44. Crowe A, Dirks C, Wenderoth MP. Biology in Bloom: Implementing Bloom’s Taxonomy to Enhance Student Learning in Biology. LSE. 2008;7: 368–381. doi:10.1187/cbe.08-050024

45. Larsen TM, Endo BH, Yee AT, Do T, Lo SM. Probing Internal Assumptions of the Revise Bloom’s Taxonomy. LSE. 2022;21: ar66. doi:10.1187/cbe.20-08-0170

46. Goedhart J. PlotsOfDifferences–a web app for the quantitative comparison of unpaired data. BioRxiv. 2019; 578575.

47. Nuzzo RL. Randomization test: An alternative analysis for the difference of two means. PM&R. 2017;9: 306–310.

48. Bates DM. lme4: Mixed-effects modeling with R. Springer New York; 2010.

49. Kuznetsova A, Brockhoff PB, Christensen RH. lmerTest package: tests in linear mixed effects models. Journal of statistical software. 2017;82: 1–26.

50. Casagrand J, Semsar K. Redesigning a course to help students achieve higher-order cognitive thinking skills: from goals and mechanics to student outcomes. Adv Physiol Educ. 2017;41: 194–202. doi:10.1152/advan.00102.2016

51. Lo S, Baiduc R, Swarat S, Drane D, Light G. When active learning is not active learning: How conceptions of teaching inform implementations of active-learning approaches. Baltimore, MD; 2016.

52. Elliott ER, Reason RD, Coffman CR, Gangloff EJ, Raker JR, Powell-Coffman JA, et al. Improved student learning through a faculty learning community: How faculty collaboration transformed a large-enrollment course from lecture to student centered. CBE—Life Sciences Education. 2016;15: ar22.

53. Stains M, Vickrey T. Fidelity of Implementation: An Overlooked Yet Critical Construct t Establish Effectiveness of Evidence-Based Instructional Practices. Allen D, editor. LSE. 2017;16: rm1. doi:10.1187/cbe.16-03-0113

54. Henderson C, Beach A, Finkelstein N. Facilitating change in undergraduate STEM instructional practices: An analytic review of the literature. J Res Sci Teach. 2011;48: 952–984. doi:10.1002/tea.20439

55. Smith MK, Jones FHM, Gilbert SL, Wieman CE. The Classroom Observation Protocol for Undergraduate STEM (COPUS): A New Instrument to Characterize University STEM Classroom Practices. Dolan EL, editor. LSE. 2013;12: 618–627. doi:10.1187/cbe.13-08-0154

56. Owens MT, Seidel SB, Wong M, Bejines TE, Lietz S, Perez JR, et al. Classroom sound can be used to classify teaching practices in college science courses. Proceedings of the National Academy of Sciences. 2017;114: 3085–3090. doi:10.1073/pnas.1618693114

57. Eddy SL, Converse M, Wenderoth MP. PORTAAL: A Classroom Observation Tool Assessing Evidence-Based Teaching Practices for Active Learning in Large Science, Technology, Engineering, and Mathematics Classes. CBE Life Sci Educ. 2015;14: ar23. doi:10.1187/cbe.14-06-0095

58. McConnell M, Boyer J, Montplaisir LM, Arneson JB, Harding RLS, Farlow B, et al. Interpret with Caution: COPUS Instructional Styles May Not Differ in Terms of Practice That Support Student Learning. Andrews TC, editor. LSE. 2021;20: ar26. doi:10.1187/cbe.20-09-0218

59. Wieman C. Response to “Interpret with Caution: COPUS Instructional Styles May Not Differ in Terms of Practices That Support Student Learning,” by Melody McConnell, Jeffrey Boyer, Lisa M. Montplaisir, Jessie B. Arneson, Rachel L. S. Harding, Brian Farlow and Erika G. Offerdahl. LSE. 2021;20: le1. doi:10.1187/cbe.21-05-0126

60. Crouch CH, Mazur E. Peer Instruction: Ten years of experience and results. American Journal of Physics. 2001;69: 970–977. doi:10.1119/1.1374249

61. Linton DL, Farmer JK, Peterson E. Is Peer Interaction Necessary for Optimal Active Learning? Hewlett J, editor. LSE. 2014;13: 243–252. doi:10.1187/cbe.13-10-0201

