## Supplementary Information for "Effect of time spent on active learning on exam performance: A controlled case study on a course with different instructors but identical teaching materials"

Supplemental Table 1
**Description of class sessions and questions analyzed**

*Observations and questions analyzed*

| Teaching unit | Learning objectives | Class sessions | Clicker questions | Exam questions |
|---|---|---|---|---|
| 1 | 7 | 3 | 19 | 43 |
| 2 | 3 | 3 | 18 | 19 |
| 3 | 4 | 3 | 18 | 8 |

Supplemental Figure 1
**Fraction of Active Learning Class Time**

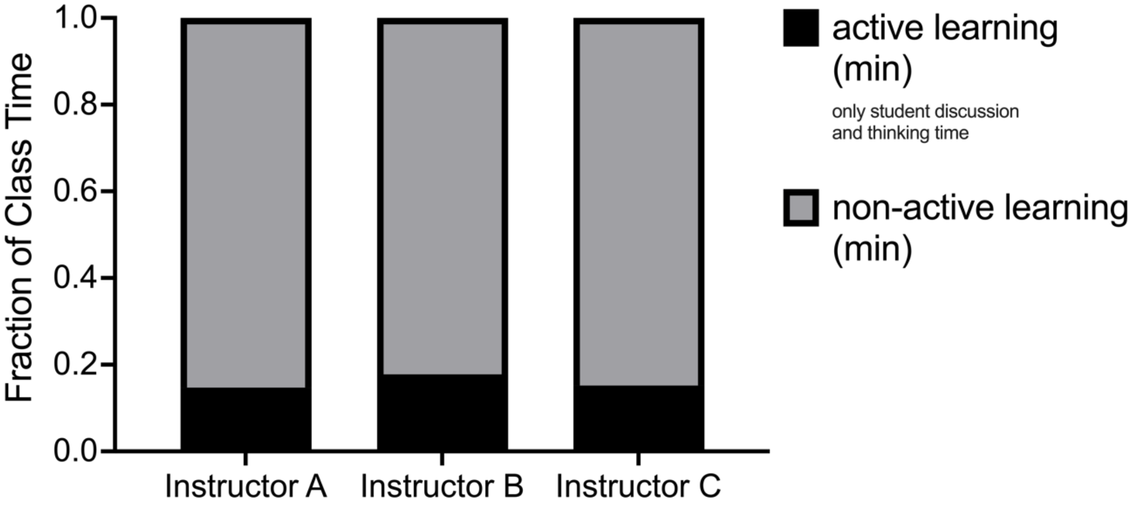

Supplemental Figure 2

**Concept Inventory Scores Across Instructors**

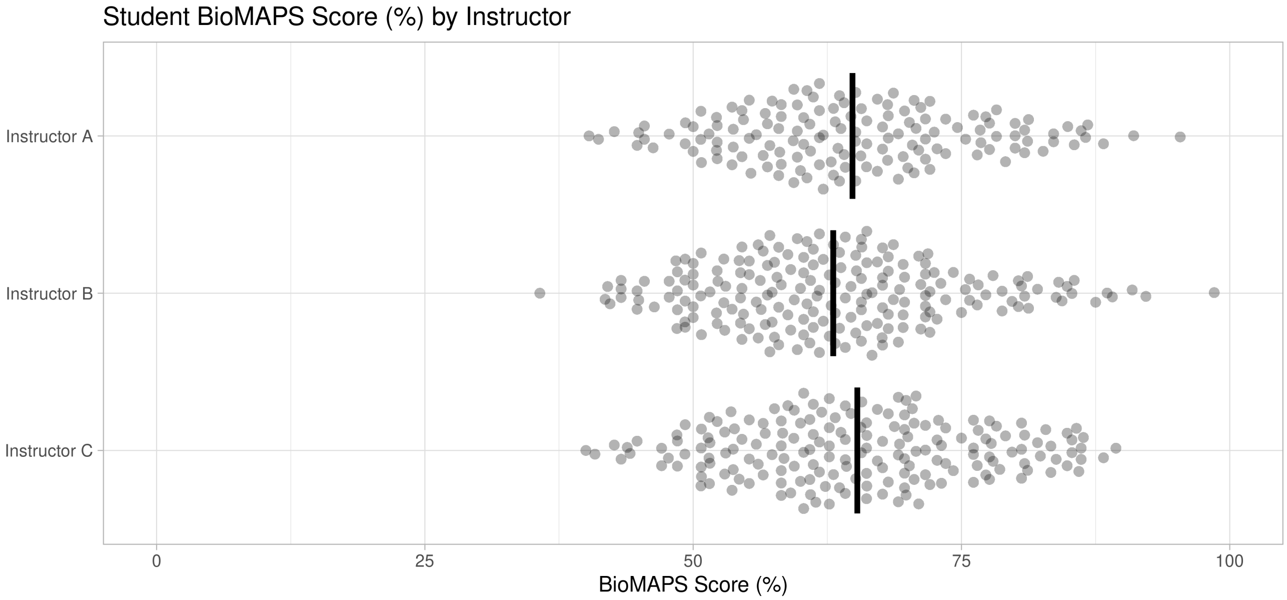

Supplemental Figure 3
**Sample Questions**

**Lower-Order Clicker Question:**
Organelles (LO: Describe the function of common eukaryotic organelles: mitochondrion, chloroplast, nucleus, endoplasmic reticulum, Golgi apparatus, lysosome, and vacuole) (Objective: 2A)

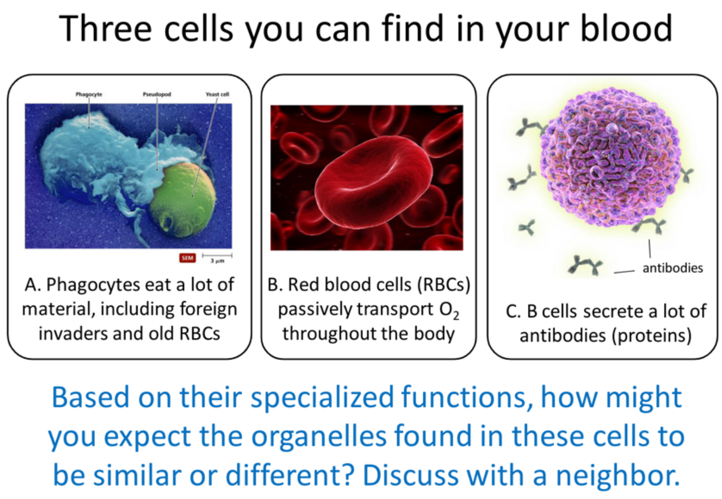


**Lower-Order Exam Question:**
(Midterm 1)

[Questions 25-26] You are studying a collection of eukaryotic cell samples in the lab. Unfortunately, all of the labels have fallen off of the samples! Each question below represents one of the cell samples you are studying. Use the answer choices [A, B, C, D, or E] to designate which organelle would be the most helpful for correctly distinguishing each sample from the other samples based on the known functions of your cells. Answer choices may be used more than once or not at all.

A. Lysosome
B. Golgi Apparatus
C. Vacuole
D. Chloroplasts
E. Mitochondria

25. Human stomach cells that secrete lots of digestive enzymes.
26. Muscle cells taken from an Olympic runner.

**Higher-Order Clicker Question:**
Epigenetics (LO: Discuss how histones are chemically modified to increase or decrease the expression of a gene) (Objective: 7B)

Mark A for True or B for False:

The gel shown below could explain the results on an RNA and Protein gel if there was no hydrolysis of ATP associated with the SWI/SNF nucleosome remodeling complex.

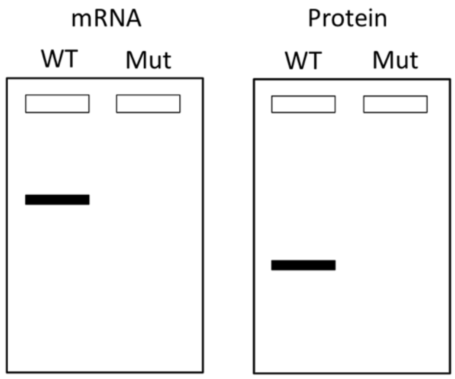


**Higher-Order Clicker Question:**
(Final)

[Questions 63-66] You are studying a human gene found on Chromosome 9. Each of the questions below describes a different mutation that could affect the structure and/or expression of this gene. Determine which Northern Blot gel below [A, B, C, D, or E] you would be <u>most likely</u> to observe as a consequence of each mutation. On each gel, "WT" is the mature mRNA produced by normal wild type cells and "MUT" is the mature mRNA produced by mutated cells. Answers may be used more than once or not at all.

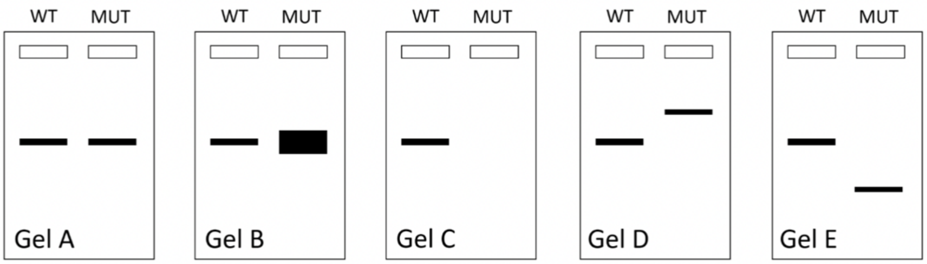


63. Gel that would result from a mutation that increases methylation of a CpG island in the promoter of this gene. **C**
64. Gel that would result from a mutation that inactivates the SWI/SNF (nucleosome remodeling) complex. **C**
65. Gel that would result from a nonsense mutation in this gene. **A**
66. Gel that would result from a mutation that inactivates the histone deacetylases (HDACs) that regulate expression of this gene. **B**

Supplemental Table 2

**Percentage of Higher-order Questions Analyzed**

| **Clicker/Exam** | **3A clicker** | **5B clicker** | **7B clicker** | **2A clicker** | **4B clicker** | **6B clicker** | **3A midterm1** | **5B midterm2** | **3A final** | **5B final** | **7B final** | **midterm1 <50** | **midterm2 <50** | **final <50** |
|---|---|---|---|---|---|---|---|---|---|---|---|---|---|---|
| **Total Questions Analyzed** | 17 | 15 | 15 | 5 | 9 | 6 | 25 | 15 | 8 | 1 | 9 | 3 | 2 | 5 |
| **Higher Order Questions** | 0 | 3 | 3 | 1 | 2 | 3 | 7 | 7 | 2 | 0 | 4 | 1 | 1 | 0 |
| **% Higher** | 0.00% | 20.00% | 20.00% | 20.00% | 22.22% | 50.00% | 28.00% | 46.67% | 25.00% | 0.00% | 44.44% | 33.33% | 50.00% | 0.00% |

Supplemental Figure 4
**Coding Guidelines**

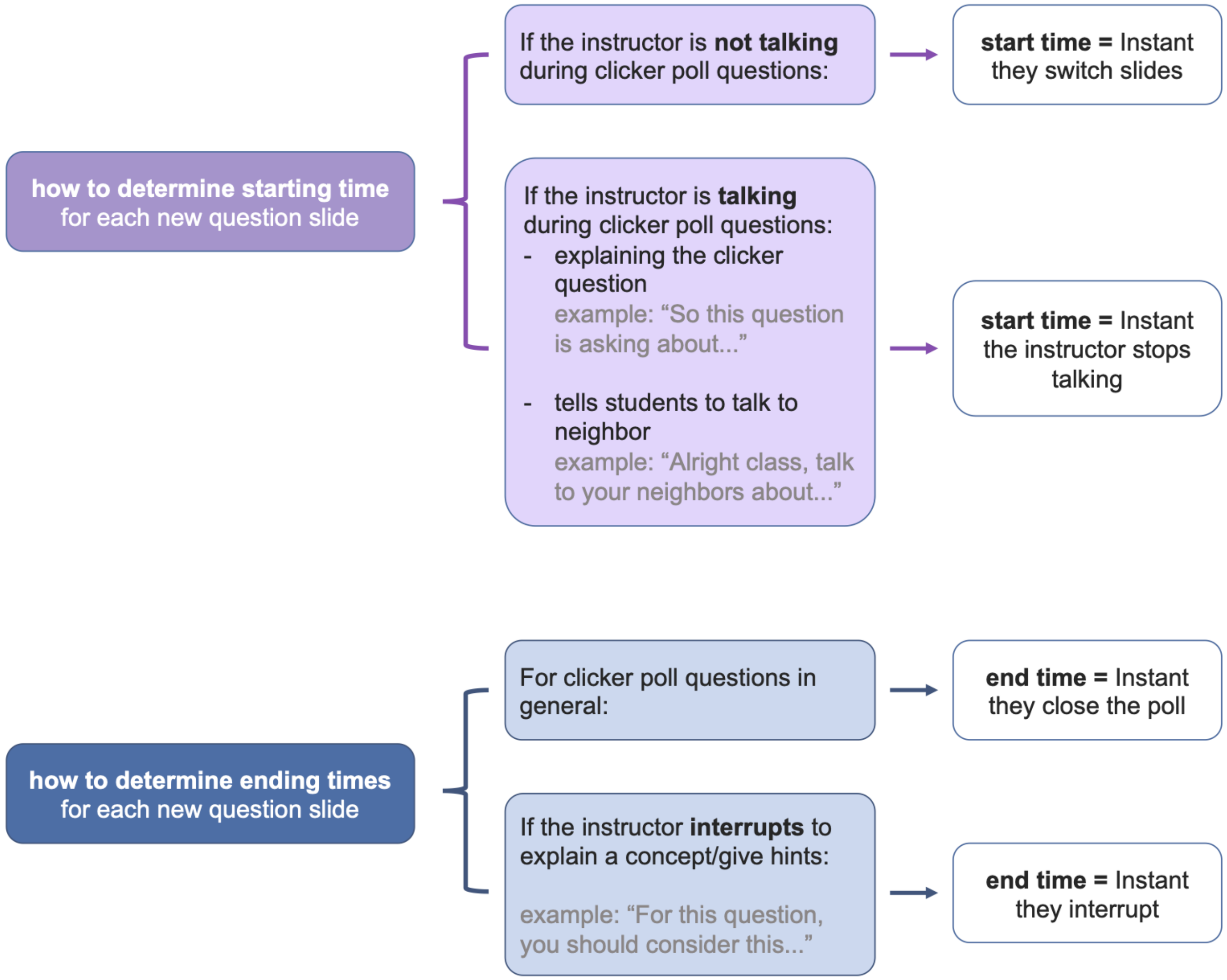
